# Post-selection Inference Following Aggregate Level Hypothesis Testing in Large Scale Genomic Data

**DOI:** 10.1101/058404

**Authors:** Ruth Heller, Nilanjan Chatterjee, Abba Krieger, Jianxin Shi

## Abstract

In many genomic applications, hypotheses tests are performed by aggregating test-statistics across units within naturally defined classes for powerful identification of signals. Following class-level testing, it is naturally of interest to identify the lower level units which contain true signals. Testing the individual units within a class without taking into account the fact that the class was selected using an aggregate-level test-statistic, will produce biased inference. We develop a hypothesis testing framework that guarantees control for false positive rates conditional on the fact that the class was selected. Specifically, we develop procedures for calculating unit level p-values that allows rejection of null hypotheses controlling for two types of conditional error rates, one relating to family wise rate and the other relating to false discovery rate. We use simulation studies to illustrate validity and power of the proposed procedure in comparison to several possible alternatives. We illustrate the power of the method in a natural application involving whole-genome expression quantitative trait loci (eQTL) analysis across 17 tissue types using data from The Cancer Genome Atlas (TCGA) Project.

## 1 Introduction

In large scale analysis of genomic and genetic data, it is common to perform tests for hypotheses at an aggregate level over ”classes” of multiple related units. In analysis of genome-wide association studies, for example, testing for genetic associations may be performed at a gene-, region- or pathway- level by aggregating association statistics over multiple genetic markers (Yu et al., 2009; Wu et al., 2010a,b). Similarly, for identification of susceptibility markers with pleiotropic effects, association tests for each individual genetic marker may be performed by aggregating association test-statistics over multiple related phenotypes (Bhattacharjee et al., 2012; Peterson et al., 2015). Recently, in microbiome-profiling studies, association tests for identifying taxa were performed by aggregating association test-statistics over multiple traits (Hua et al., 2016). When multiple signals are expected within classes of related units, use of suitable aggregate-level test-statistics can improve the power of discovery of the signal harboring classes, compared to one unit at a time analysis.

In using aggregate level test-statistics for large scale hypotheses testing, much research have focused on development of test-statistics that can combine evidence of signals from multiple units in the most powerful way. For any given test-statistics, standard multiple testing adjustment methods are typically applied to adjust for the number of classes, which could be potentially large, over which the analyses may be performed. An important area of research that has received less attention involves how to perform proper post-selection inference following the aggregate level hypothesis testing to isolate individual units that contain signal within the higher-level classes. For example, if a genetic marker is identified to be statistically significant for aggregate-level association in pleiotropic analysis across multiple phenotypes, it is then very natural to perform follow-up analysis to identify which underlying phenotypes contain true signals for associations.

Clearly, a naive approach that ignores the effect of class selection on the second-stage of individual units selection will produce biased inference even if multiple testing adjustment methods were used to adjust for the number of units within each selected class.

In this manuscript, we study the problem of post-selection inference following aggregate level analysis as a two-stage hypothesis testing problem. The data of interest, the unit-level test-statistics, can be organized in an *m* × *n* matrix, where *m* is the number of classes for which inference is of interest, and *n* is the number of units within each class. For example, in GWAS of different phenotypes the data matrix for analysis has rows indexed by the SNPs, and columns indexed by the phenotypes. The first goal is to select the rows (SNPs) that have signal in at least one column (phenotype), and the post-selection goal can be to identify which columns have signal within each of the chosen rows.

Approaches to various error rate controls have been proposed in the past for multi-stage or hierarchical hypothesis testing. Possibly most relevant of them in the context of applications described above is a procedure proposed by Benjamini and Bogomolov (2014) which control false positives on the average over the selected rows. The method was recently applied to pleiotropic analysis of GWAS and single tissue eQTL data (Peterson et al., 2015, 2016), a type of application that we also use for illustration in this manuscript. Barber and Ramdas (2015) suggested controlling the FDR at an aggregate level of rows or columns, in addition to controlling the FDR across all individual hypotheses. Liu et al. (2016) suggested controlling a posterior measure of false discoveries within each selected row, in addition to overall. Yekutieli (2008) suggested controlling different FDR-types on trees of hypotheses, assuming independence between the stages of the hierarchy. Some other strategies have been tailored to special application fields, such as gene expression analyses (Yekutieli et al.,, 2006; Li and Ghosh, 2014), electrencephalography research (Singh and Phillips, 2010), and functional magnetic resonance imaging (Schildknecht et al., 2015). Conditioning on the selection event has been suggested in novel works on post-model selection (Fithian et al., 2015; Lee et al., 2016), as well as for spatial signals in Benjamini and Heller (2007).

For the selection of columns in the second-stage of above hypothesis testing framework, we propose developing multiple testing procedures that guarantees control of false positive rates conditional on the fact that the row is selected. As a researcher may often conduct different experiments for each selected row for follow-up studies, control of false positive at the level of each row may be desirable rather than controlling average rates over all selected rows as has been considered for this problem by Peterson et al. (2015). It has been noted earlier (Benjamini and Bogomolov, 2014) that development of a general procedure for controlling such error rates could be difficult. In the setting where the test-statistics can be assumed to be independent across columns, we propose general procedures that are theoretically shown to control family-wise error rates and false discovery rate at the level of each row conditional on selection. We then show through simulation studies that for a commonly used aggregation test-statistic, the proposed procedure not only provides stronger type-I error rate control but it also has a major power advantage when compared to the general procedure proposed by Benjamini and Bogomolov (2014). An application of the methods for analysis of SNP markers predictive of gene expression level across multiple tissues using data from the The Cancer Genome Atlas (TCGA) project provided further empirical validation of substantially higher power of the proposed method compared to alternatives. These results provide encouraging direction for further research in developing more powerful procedures for two-stage hypothesis testing in more general frameworks.

In section 2, we set up the proposed hypothesis testing framework more formally and define the criterion of type-I error that we propose to control. In section 3, we present a procedure for controlling the proposed type-I error criterion and provide theoretical results supporting its validity. In section 4, we conduct simulation studies to evaluate type-I error rates and power for the proposed method compared to a naive approach and to the procedure proposed by Benjamini and Bogomolov (2014). In section 5, we describe results from analysis of TCGA study. In Section 6, we conclude with a discussion about future directions.

## 2 Notation and Goal

Let *H_ij_, j* = 1,…,*n* be the family of *n* null hypotheses for the ith row, and let *P_ij_, j* = 1,…, *n* be their corresponding *p*-values, for *i* = 1,…, *m*. The global null hypothesis for row *i*, 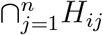, is the hypothesis that all null hypotheses *Hij*, *j* = 1,…,*n* are true. Let *P_iG_* be the global null *p*-value for row *i*, i.e., a valid *p*-value of the global null hypothesis 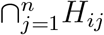. *P_iG_* is based on the *p*-values *p_ij_*, *j* = 1,…, *n*, or on the corresponding unit-level test-statistics. For example, it can be computed as in equation (3.1), or equation (5.1), or as suggested in Remark 3.1. Denoting the observed *p*-values by lower-case letters, our data matrix for analysis is:

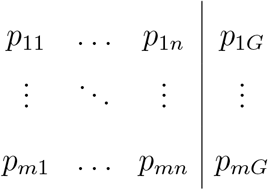

The first stage analysis selects rows based on the global null *p*-values *p*_1*G*_,…,*p_mG_*, to answer the question of which rows are promising, i.e., show evidence that there is signal in at least one column. Let 𝒮 ⊆ {1,…, *m*} be the set of selected rows. For example, when the rows are the SNPs and the columns are different studies, the selected rows are often those that achieve genome-wide significance with FWER control, so 𝒮 = {*i*: *p_iG_* ≤ *α*/*m*}.

The second stage analysis is based on the individual *p*-values {*p_ij_*: *j* = 1,…, *n*, *i* ∈ 𝒮 }, to answer questions such as which columns within selected rows contain signal.

Let *V_i_* and *R_i_* be the number of false and total rejections for row *i*, respectively. Our first error measure, which henceforth is referred to as the conditional FWER for a selected row, is the probability of at least one erroneous rejection within the row, conditional on it being selected,

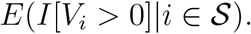

Our second error measure, which henceforth is referred to as the conditional FDR for a selected row, is the expected fraction of erroneous rejections among the rejections within the row, conditional on it being selected,

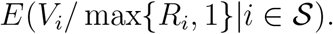

If *n* = 1, our error measures coincide with the selective type I error rate, i.e. the error rate of a test given that it was performed, as suggested by Fithian et al. (2015).

Our goal is to suggest valid multiple testing procedures for controlling the conditional FWER/FDR for the selected rows.

## 3 Valid inference within a selected row

In this section we provide procedures for controlling the conditional FWER/FDR for a selected row. The valid inference we provide relies on the assumption that the the columns are independent, as is typical in meta-analyses. However, across rows (i.e., within columns), the individual level test-statistics (or *p*-values) may be dependent.

The case where a row is selected if the aggregate test passes a predefined threshold is addressed in § 3.1. We provide procedures that control the conditional FWER/FDR for any type of dependency across rows. This type of selection is standard practice in GWAS meta-analysis, see § 5 for a rich data application.

The case where a row is selected based on a data-dependent threshold is addressed in § 3.2. We provide procedures that for independence across rows control the conditional FWER/FDR, as well as the average error rate (Benjamini and Bogomolov, 2014). We discuss conditional and average error rate control using our approach when there is dependency across rows.

### 3.1 Inference following selection of rows using a fixed cut-off on the aggregate level test-statistics

Even in the simplest setting where the row selection is based on a fixed cut-off, adjusting for selection is not obvious since the probability of the selection event depends on the unknown distribution of the non-null *p*-values within the row. In § 3.1.1 we suggest a way around this problem, which leads to valid conditional *p*-values. Although the original *p*-values were independent, the conditional *p*-values in the selected row are dependent due to the selection event. In § 3.1.2 we present multiple testing procedures based on these conditional *p*-values that control the conditional error rate.

#### 3.1.1 The conditional p-value computation

For each row *i* ∈ {1,…, m}, the *p*-values corresponding to the different columns are assumed independent. The computation of *p_iG_* can be based on different aggregates of the *p*-values *p_ij_*, *j* = 1,…,*n*, see Loughin (2004) for common choices and a review. A popular choice is Fisher’s combining method, which has been shown to have excellent power properties (see, e.g., Owen (2009) and the references within). Let f: ℛ^n^ → ℛ be a combining function for testing the global null on each row. For example, Fisher’s combining function is 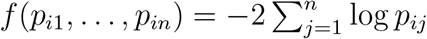. Under the global null of no signal in the row, if the *p*-values in the row are uniformly distribution, 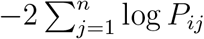 has a chi-squared distribution with *2n* degrees of freedom, 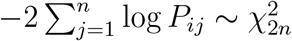, so

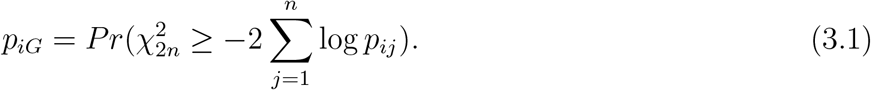

See Remark 3.1 and Section 5 for other combining functions.

Let *t* be the fixed row selection cut-off, so that row *i* is selected if f (*p_i1_*,…,*p_in_*) ≥ *t*. We require that the combining method be non-increasing in each of its arguments, when the other *n* − 1 values are held fixed. This is a reasonable requirement since if the individual *p*-values decrease, the selection of the row should become easier, so if f (p_i1_,…, p_in_) ≥ t and 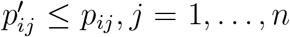, then 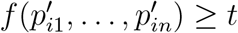. For simplicity, we shall assume that the *p*-values are continuous, with a uniform null distribution, and therefore that *f* is continuous.

Since we assume we are dealing with a selected row based on the global test, we will omit the index *i* for row from hereon to simplify the notation. The *n* independent *p*-values for a selected row are therefore denoted by *P_1_*,…, *P_n_*, and their observed values are *p_1_*,…*, p_n_*. We observed that we rejected the global null which means that the *p*-values satisfy f (p_1_,…, p_n_) ≥ t.

We let *b_j_* satisfy

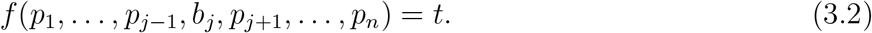

Clearly, b_j_ ≥ p_j_. If no such b_j_ exists, that is f (*p*_1_,…,*p*_*j*−1_, 1,*p*_*j*+1_,…,*p_n_*) ≥ *t*, let *b_j_* = 1.

The aim is to provide a test for each of the n hypotheses having observed that the global null was rejected. To this end, we adjust the *p*-values to

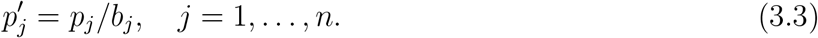

These are valid conditional *p*-values. To see this, note that *P_j_* conditional on the row being selected and on the remaining *p*-values is truncated at *b_j_* ∈ (0,1]. So if unconditionally *P_j_* is uniformly distributed, *P_j_* ~ *U*(0,1), then conditionally on being selected and on the remaining *p*-values it has a uniform distribution between zero and b_j_,

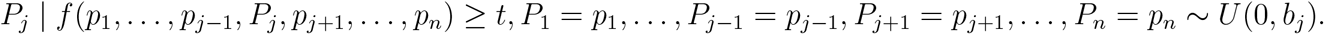

For Fisher’s combining function, *f*(*p*_1_,…,*p_n_*) ≥ *t* is equivalent to 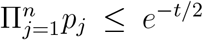. Therefore, 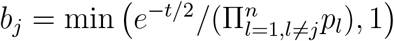, and the conditional *p*-values are:

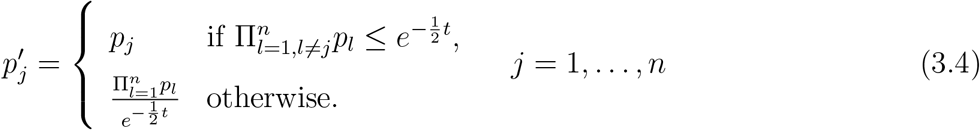

The magnitude of the inflation in the post-selection *p*-values, *p*_*j*_^′^− *p_j_*, depends on how the signal is distributed in the row. Using Fisher’s combining method, there can be no inflation (i.e., no cost for selection!) if the signal is strong in at least two columns. This is a direct result of the conditional *p*-value computation (3.4), where *p_j_*^′^ = *p_j_* if 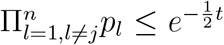. Note that even though it is enough to have one strong signal for the selection of the row, at least two columns need to contain strong signals for all n conditional *p*-values to coincide with the original *p*-values. Therefore, computing the conditional *p*-values as suggested in (3.3) will typically lead to smaller conditional *p*-values whenever signals are present in more than one column, than the use of the following probabilities, which are computed under the global null:

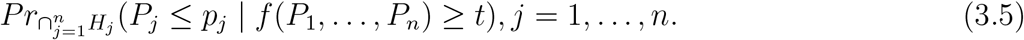

The computation in (3.5) uses conservatively the uniform distribution for the *p*-values, *P_k_ ~ U* (0,1) for *k* = 1,…, *n*, and the resulting probabilities can be substantially larger than the original *p*-values: for Fisher’s combining method, if *t* = χ_1−α/m,2n_ and 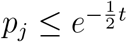, the value will be 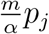.

##### Remark 3.1

(Other *p*-values combining methods). *Conceptually, our approach for computing the conditional p-values can be applied to many different tests of the global null. The complexity of the conditional p-value computation in (3.3) can vary greatly between combining methods. For one-sided tests, the computation is straightforward using Stouffer’s combining method, f*(*p*_1_,…,*p_n_*) = 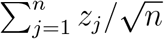, *where z_j_* = Φ^−1^(1 — *p_j_*), *which has a standard normal distribution under the global null. So the global p-value is* 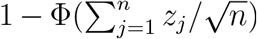. *If the row is selected when* 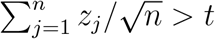, *then the conditional p-values are:*

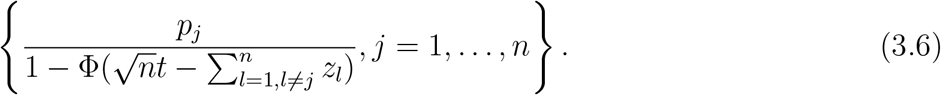

*The computation of the conditional p-values is more complex using the test of Bhattacharjee et al. (2012) for the global null, called ASSET. Their test is based on identifying the set S_max_ that has the largest weighted Stouffer test statistic, and the significance at the aggregate row level is based on the maximal test statistic*, 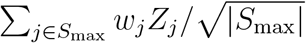 *Bhattacharjee et al. (2012) showed that their global test can be more powerful than a test statistic that aggregates all the information in the row without subset selection*.

##### Remark 3.2

(The ranking of the original and conditional *p*-values). *Using Fisher’s or Stouffer’s combining method, it is easy to show that the ranking of the p-values remains unchanged, i.e., if p_1_ ≤ … ≤ p_n_ then p_1_^′^ ≤ … ≤ p_n_^′^. Other combining methods may lack this desired property. For example, if the global null p-value is based on Bonferroni, i.e.*, *p_G_* = *n* × min{*p*_1_,…,*p_n_* }, *the ranking is changed following selection. To see this, suppose that the row is selected if p_G_ < α/m, with p_i_ = α/(m × n) but pj ∈ (α/m, 1), j = 2,…,n. Then b_1_ is the value that satisfies n × min{b_1_,p_2_,…, p_n_ } = α/m, i.e., b_1_ = α/(m × n), so p_1_^′^ = 1, but p_j_^′^ = p_j_* < 1.

#### 3.1.2 >Valid procedures for conditional FWER/FDR control

Applying a valid FWER/FDR controlling procedure on {*p_j_*^′^: *j*=1,…,*n*} will guarantee control of the conditional FWER/FDR for a selected row. The choice of FWER/FDR controlling procedure should be guided by the observation that the *n* conditional *p*-values in the row are dependent, despite the fact that the original *p*-values were independent across columns. The conditional FWER for a selected row will be controlled if we use Bonferroni-Holm on the conditional *p*-values within each selected row, since the Bonferroni-Holm procedure guarantees FWER control for any dependency between the *p*-values. For conditional FDR control, we would like to use the Benjamini-Hochberg (BH) procedure (Benjamini and Hochberg, 1995) within each selected row at level *α*. The next theorem shows that for the dependency among the *p*-values induced by the conditioning step, the BH procedure is indeed valid.

##### Theorem 3.1.

*Let P_1_,…,P_n_ be independent p-values, each with a uniform null distribution. If f(p_1_,…,p_n_) > t, then the BH procedure at level α on p_1_^′^,…,p_n_^′^ controls the conditional FDR at level* 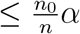, *where n_0_ is the number of true null hypotheses*.

*Proof*. Without loss of generality, relabel the columns so that the first *n*_0_ columns have a true null hypothesis. We make use of the representation of FDR from Benjamini and Yekutieli (2001), so the conditional FDR is

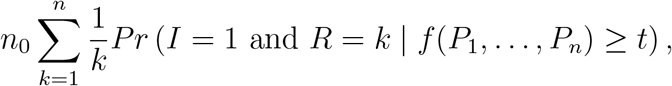

where *I* = 1 if *H_1_* (the first hypothesis), which we assume to be null, is rejected, and *R* is the number of rejected hypotheses. We condition on *p_2_,…,p_n_* so that it is sufficient to show that

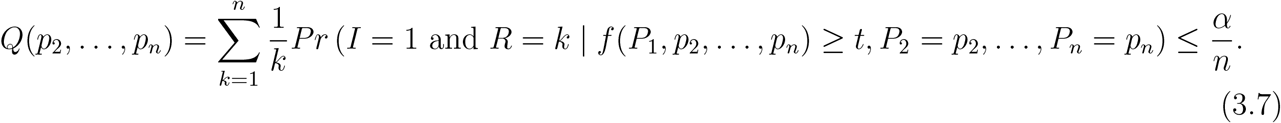

The only random quantity in *Q* is *P_1_^′^* = *P_1_/b_1_*, and it is uniformly distributed between 0 and 1 given the selection event. As *p*_1_^′^ increases *b_2_*,…, *b_n_* will be non-increasing so there must be 0 = *a_0_ < a_1_ < … < a_L_* = 1 so that *R*(*p_1_^′^*) = *k_l_* for *a*_*l*−1_ ≤ *p_l_^′^ ≤ a_l_*,*l* = 1,…,*L*, where *k_1_ > k_2_ > … > k_L_*. Since we need *I* = 1, or p_1_^′^ < R(p_1_^′^)α/n, there exists *t* such that

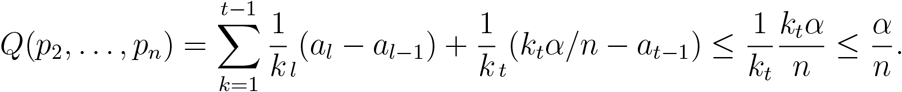

Interestingly, for Fisher’s selection rule the conditional FDR is exactly n_0_*α/n*.

##### Corollary 3.1.

*Let P_1_,…,P_n_ be independent p-values, each with a uniform null distribution. If 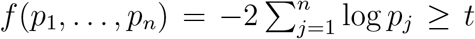, then the conditional FDR of the BH procedure at level α on p_1_^′^,…,p_n_^′^ is equal to 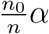*.

See Appendix A for the proof.

##### Remark 3.3

(Estimation of the fraction of nulls in selected rows). *Theorem 3.1 shows that the FDR of the level-α BH procedure applied on the conditional p-values is bounded by n_0_a in a selected row, where π_0_ = n_0_/n. When n is large enough (say 30 or more), it may be useful to estimate n_0_ and incorporate the estimate in the multiple testing procedure to gain power. The gain in power can be substantial if π_0_ is far less than one, as appears to be the case in the application considered in Section 5. Schweder and Spjotvoll (1982) proposed estimating π_0_ by* 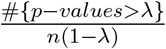, *where λ ∈ (0,1). The slightly inflated plug-in estimator* 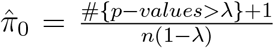 *has been incorporated into multiple testing procedures in recent years. For independent p-values, Storey (2003) showed that the BH procedure at level α/(π̂_0_) controls the FDR at level α. Benjamini et al. (2006) suggested another estimator, and noted that the BH procedure which incorporates the plug-in estimator with λ = 0.5 is sensitive to deviations from the assumption of independence, and its FDR level may be inflated above the nominal level under dependency. Blanchard and Roquain (2009) provided extensive simulations that show that the BH procedure which incorporates the plug-in estimator with λ = α is robust to deviations from the assumption of independence, at the price of being slightly more conservative. We need to investigate which estimator will result in an adaptive BH procedure that controls the conditional FDR. Also, a confidence statement about π_0_ within selected rows may be of interest in itself, in providing an upper bound on the fraction of nulls (or a lower bound on the fraction of columns with signal) within each selected row*.

### 3.2 Inference following data adaptive row-selection rules

In § 3.2.1 we show that our theoretical results carry over to more general row selection rules when the test statistics within each column are independent. In § 3.2.2 we show the connection with the average error rate (Benjamini and Bogomolov, 2014). In § 3.2.3 we discuss conditional and average error rate control using our approach when the test statistics within each column are dependent.

#### 3.2.1 Valid inference under row independence

The results in § 3.1.2 hold if t_i_ does not depend on *P_i1_,…, P_in_*. The independence between the threshold and the ith row *p*-values is clearly satisfied in the setting of § 3.1, where the thresholds is fixed given m and n (or more generally, given the number of non-missing observations in row *i*, say *n_i_*). For example, for GWAS, the threshold is often chosen so that the FWER on the m global tests is controlled at the desired nominal level α. If rows are selected by Bonferroni, then the selection threshold for Fisher’s combining method is t = *χ_1−α_/m,_2n_*, i.e., the (1 − *α/m)* quantile of 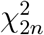

If the *p*-values across rows are independent, more general data-adaptive thresholds can also lead to valid inference within selected rows. This is clearly so if the threshold for selection depends on all *p*-values except the *p*-values in the selected row, i.e., row i is selected if f (*p*_*i*1_,…, *p_in_*) ≥ *t*(*p*_−*i*_), where p_−i_ = {p_kl_: *k* = 1,…, *i* − 1, *i* + 1,…, *m*, *l* = 1,…, *n*}. If we compute the conditional *p*-values for selected row *i* as in (3.3) with *t* = *t*(*p*_−*i*_), the post-selection inference using Bonferroni-Holm or BH on the conditional *p*-values at level *α* will control, respectively, the conditional FWER or conditional FDR at level *α*.

For example, if the selected rows are the discoveries form a BH procedure at level q on the global null *p*-values, as suggested by Peterson et al. (2015), the conditional post-selection inference will control the conditional error. Specifics follow. Relabelling a selected row as row number one, and the remaining rows in order of their global null *p*-values, i.e., *p*_2*G*_ ≤ … ≤ *p_mG_*, let 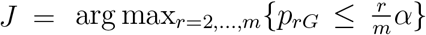. If no such r exists then *J* =1. Row 1 is selected if *p_iG_* ≤ *Jq/m*. Clearly, the threshold for selection Jq/m is independent of *P*_11_,…, *P*_1*n*_ if the *p*-values across rows are independent. Therefore, the conditional *p*-values for each selected row are computed by treating the selection threshold as fixed. For Fisher’s combining function, the selection threshold for a selected row is 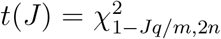, and the conditional *p*-values for each selected row are computed by the formulas in (3.4) with *t* = *t*(*J*).

The class of simple selection rules, which includes row-selection by the BH procedure or by Bonferroni-Holm on the global null *p*-values, leads to valid conditional inference within selected rows.

##### Definition 3.1

(Definition 1 in Benjamini and Bogomolov (2014)). *A selection rule is called simple if for each selected row, when the p-values not belonging to that row are fixed and the p-values in that row can change as long as the row is selected, the number of selected rows remains unchanged*.

For each *i* ∈ 𝒮, let *P*^(*i*)^ = (P_1G_, …, *P*(_*i*−1_)*_G_*, *P*(*_i_*+_1_)*_G_*,…, *P_mG_*) be the vector of *p*-values excluding *P_iG_*.

##### Theorem 3.2.

*Assume that rows are selected by a simple selection rule such that row i is selected if f(p_i1_,…,p_in_) ≥ t(|𝒮|). If the p-values across rows are independent, then if we compute the conditional p-values for selected row i as in (3.3) with t = t(|𝒮|)*:

1. *E(I[V_i_* > 0]|*i ∈ 𝒮,P*^(*i*)^) ≤ *α if we use the Bonferroni-Holm procedure on the conditional p-values*.
2. *E*(*V_i_*/max {*R_i_*, 1}|*i* ∈ 𝒮, *P*^(*i*)^) ≤ *α if we use the BH procedure on the conditional p-values*.

See Appendix B for a proof. Let *C_i_* be the (unobserved) random variable whose conditional expectation is the desired error rate, so *C_i_* = *I* [*V_i_* > 0] for FWER control, and *C_i_* = *V_i_*/*max*(*R_i_*, 1) for FDR control. It follows from Theorem 3.2 that the conditional error is controlled, since *E*(*C_i_*|*i* ∈ 𝒮) = *E*{[*E*(*C_i_*|*i* ∈ 𝒮,*P*^(*i*)^)]|*i* ∈ 𝒮}.

#### 3.2.2 Relation to the approach of Benjamini and Bogomolov (2014)

Benjamini and Bogomolov (2014) considered selective inference on families of hypotheses, which are defined by rows in our paper. They showed that applying a Bonferroni procedure in each selected row may result in a highly inflated conditional FWER when the selection is based on within-row *p*-values. Moreover, they noted that the goal of conditional control for any combination of selection rule and testing procedure and for any configuration of true null hypotheses is difficult to achieve. Indeed, our procedures for conditional control rely on the ability to compute the conditional *p*-values, which requires well-defined selection rules and independence across the *p*-values within the rows. Benjamini and Bogomolov (2014) considered a different error measure addressing selective inference, which can be controlled under more general conditions. Specifically, they suggested to control an expected average error measure over the selected rows. While they considered a general class of error rates, we focus on two special cases. The control of the FWER on the average, 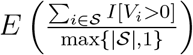, and the control of the FDR on the average, 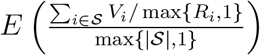

Controlling the FWER/FDR on the average is useful when we require control over false positives on the combined set of discoveries across selected rows. When we require control over false positives within each selected row separately, the conditional FWER/FDR is more appropriate.

Conditional error control can guarantee average error control, as stated in the next theorem.

##### Theorem 3.3.

*Assume row *i* is selected if *f* (*p*_*i*1_,…,p_in_) ≥ t(|𝒮|)*. *If E*(*C_i_*|*i* ∈ *𝒮,P*^(*i*)^) ≤ *α, then* 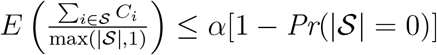.

See Appendix C for the proof which is straightforward. We know the condition is satisfied if the global null *p*-values are independent and the selection rule is simple from Theorem 3.2.

Benjamini and Bogomolov (2014) showed that by applying an FWER/FDR controlling procedure within each row at level |𝒮|α/*m*, if the set 𝒮 is selected by a simple selection rule, then the FWER/FDR on the average is controlled at level *α*. However, the conditional FWER/FDR for a selected row may exceed *α* in some of the rows as our simulations show. Comparing the conditional approach to the approach of Benjamini and Bogomolov (2014), we can generally say that the conditional approach is most sensitive to the number of non-null columns in the row, and the approach of Benjamini and Bogomolov (2014) is most sensitive to the fraction of selected rows, |𝒮|/*m*. Our simulations and real data example demonstrate that for a small fraction of selected rows, |𝒮|/*m*, the conditional approach is expected to make more discoveries than the approach of Benjamini and Bogomolov (2014). However, for sparse signal within selected rows (signal presence in only two columns or less), the approach of Benjamini and Bogomolov (2014) may make more discoveries.

#### 3.2.3 Error control following selection under row dependence

So far, we assumed that within each study, the test statistics are independent and hence the global null *p*-values are independent. We now consider the case that the studies (columns) are independent, but within each study (column), the set of all *p*-values may be dependent. This setting was considered in Benjamini and Bogomolov (2014), and although their procedures do not guarantee the control of the error rate on the average for general dependency, they prove that the error rate on the average is controlled if the *p*-values are positive regression dependent on the subset (PRDS) of true null hypotheses (Benjamini and Yekutieli, 2001).

With dependence, the global null *p*-values are no longer independent. Clearly, dependence has no effect on the conditional error guarantees of Bonferroni-Holm or the BH procedure on the conditional *p*-values, as long as the selection threshold does not depend on the global null *p*-values. In genomic applications that aim to select only rows with signal, e.g., GWAS, no false positives at the row-level are typically tolerated and a fixed selection threshold is used for FWER control on the family of global null hypotheses. For such applications, the conditional approach provides the desired theoretical guarantee over false positives within selected rows. On the other hand, the theoretical guarantee over false positives provided by the approach of Benjamini and Bogomolov (2014) is only valid if the dependency within columns is PRDS.

With a simple selection rule such as BH on the global null *p*-values, the conditional error may not be maintained. In § 4 we show empirically that for dependent column *p*-values, the conditional error is controlled, as expected, when the selection rule is fixed, but it may be inflated when the BH procedure is used for selection. However, the average error rate remained below the nominal level even with BH row selection, and it was highest when the signal was strongest.

In the Supplementary Material (SM) we provide theoretical results when the *p*-values are independent within rows, but PRDS in each column across the rows. We show that if all m rows are null rows with only true null hypotheses, the conditional approach when BH is used both for the global hypotheses at level *q* and within each row at level *α*, controls the average FDR at level ≤ *α × q* ≪ *α*. We also show that if some rows are non-null with at least one false null hypothesis, then if the *p*-values for the global null hypotheses of non-null rows is ostensibly 0, using the conditional approach when BH is used both at the global level and within each row also controls the average error at a level close to *α*.

## 4 Simulations

In order to assess the performance of different post-selection approaches, we carry out simulation studies. We have three specific aims in this simulation: (1) to examine the possible inflation in the number of false positives when failing to account for the selection (i.e., when using the original *p*-values in selected rows); (2) to compare the power of methods that properly account for selection, i.e., our novel approach that uses conditional *p*-values, and the approach of Benjamini and Bogomolov (2014); and (3) to study the effect of dependency within the columns.

We have two data generation settings. First, the row independence setting, for which all test statistics are independent. We sample the unit-level test statistics, *Z_ij_*, *i* = 1,…,*m*, *j* = 1,…,*n*, independently from the normal distribution as follows. If the null hypothesis *H_ij_* is true, the test statistic has a standard normal distribution; if the null hypothesis Hj is false, the test statistic has a normal distribution with mean *μ* ∈ {0.5,…, 7} and variance of one. The p-value is *p_ij_* = 1 − Φ(*z_ij_*), where Φ(·) is the standard normal cumulative distribution function. In each of n columns, *m* = 1000 rows are examined, and *m*_1_ = 10 rows contain signal in *n_1_* of the n columns. So we generated *n_1_* x m _1_entries in the *n* x *m* matrix of *p*-values that contained signal we aim to discover.

Second, the row-dependence setting, for which within each column of *m* hypotheses, we generate *m/B* independent blocks with within block dependency. The covariance matrix of the *B z*-scores within a block was 

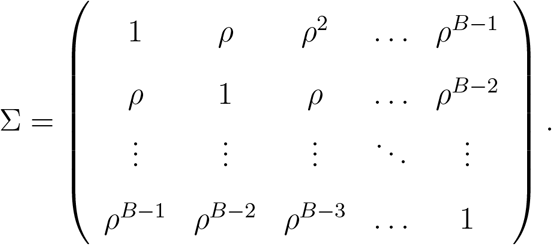

The vector of B *z*-scores within the block is multivariate normal with covariance Σ. If the block does not contain signal the mean is zero, and if it contains signal the mean is

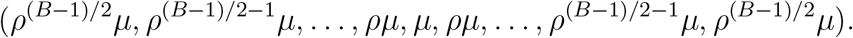

This data generation is an idealized setting for GWAS, where the SNPs are dependent due to linkage-disequilibrium, and the shape of the signal *μ* reflects the fact that the central SNP is the causal SNP and all other SNPs have signal only because of their linkage-disequilibrium with the causal SNP. In each of (*m/B*) = 100 blocks of size B = 11, one block is non-null with *μ* ∈ {0.5,…, 7} in *n_1_* of the *n* columns. So *n*_1_ × *B* entries in the *n* × *m* matrix of *p*-values contained signal we aim to discover. We varied the dependence strength ρ ∈ {0.7, 0.9, 0.99}.

The selection of rows was done by applying the Bonferroni procedure at level *α* = 0.05 or the BH procedure at level *q* ∈ {0.05, 0.2} on the global null *p*-values, computed using equation (3.1). So row *i* is selected if the hypothesis corresponding to *p_iG_* is rejected by the multiple testing procedure on {*p_iG_*: *i* = 1,…, *m*}.

We consider the following post-selection analyses using the BH procedure. The BH procedure on each selected row *i* ∈ 𝒮: at level *α* on the original *p*-values (BH-naive); at level 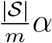 on the original *p*-values, as suggested by Benjamini and Bogomolov (2014) (BH-BB); at level *α* on the conditional *p*-values in (3.4) (BH-cond). We also considered using the Bonferroni-Holm procedure, and these results turned out to be qualitatively the same as the results using the BH procedure, see details in the SM.

To evaluate the power of the different procedures, we computed the average power, i.e., the average number of true rejections divided by the number of entries with true signal, as well as the average power for a specific row, i.e., the average number of true rejections in the row divided by *n*_1_.

### 4.1 Results for the independence setting

The results on the average and conditional FDR control show that the control of false positives was tightest using the conditional approach, since it controlled all the error measures (as expected) at the nominal 0.05 level. The approach of Benjamini and Bogomolov (2014) controlled at the nominal 0.05 level the average FDR/FWER (as expected), as well as the conditional FDR/FWER for a correctly selected row (i.e., a row that contains signal), but did not control the conditional FDR/FWER for an incorrectly selected row (i.e., a row that contains no signal). The naive approach exceeded the nominal 0.05 level in all error measures considered (as expected, since this procedure does not account for selection). Figure 1 shows the actual level of each error measures for the three post selection analyses procedures. We see that BH-naive can have a high inflation of false positives. The inflation for an incorrectly selected row was so high that it was not plotted in row 3. The fact that the inflation can be substantial clearly demonstrates that accounting for selection is necessary even when the selection criterion is very stringent. The fourth row of Table S1 in the SM shows the exact levels of the error measures for BH-naive for the setting in the last row of Figure 1 when *μ* = 4: on a correctly selected row (i.e., a row that contains signal), the conditional FDR was 0.065; on an incorrectly selected row the conditional FDR level was 0.996; and the average FDR was 0.074.

**Figure 1:**
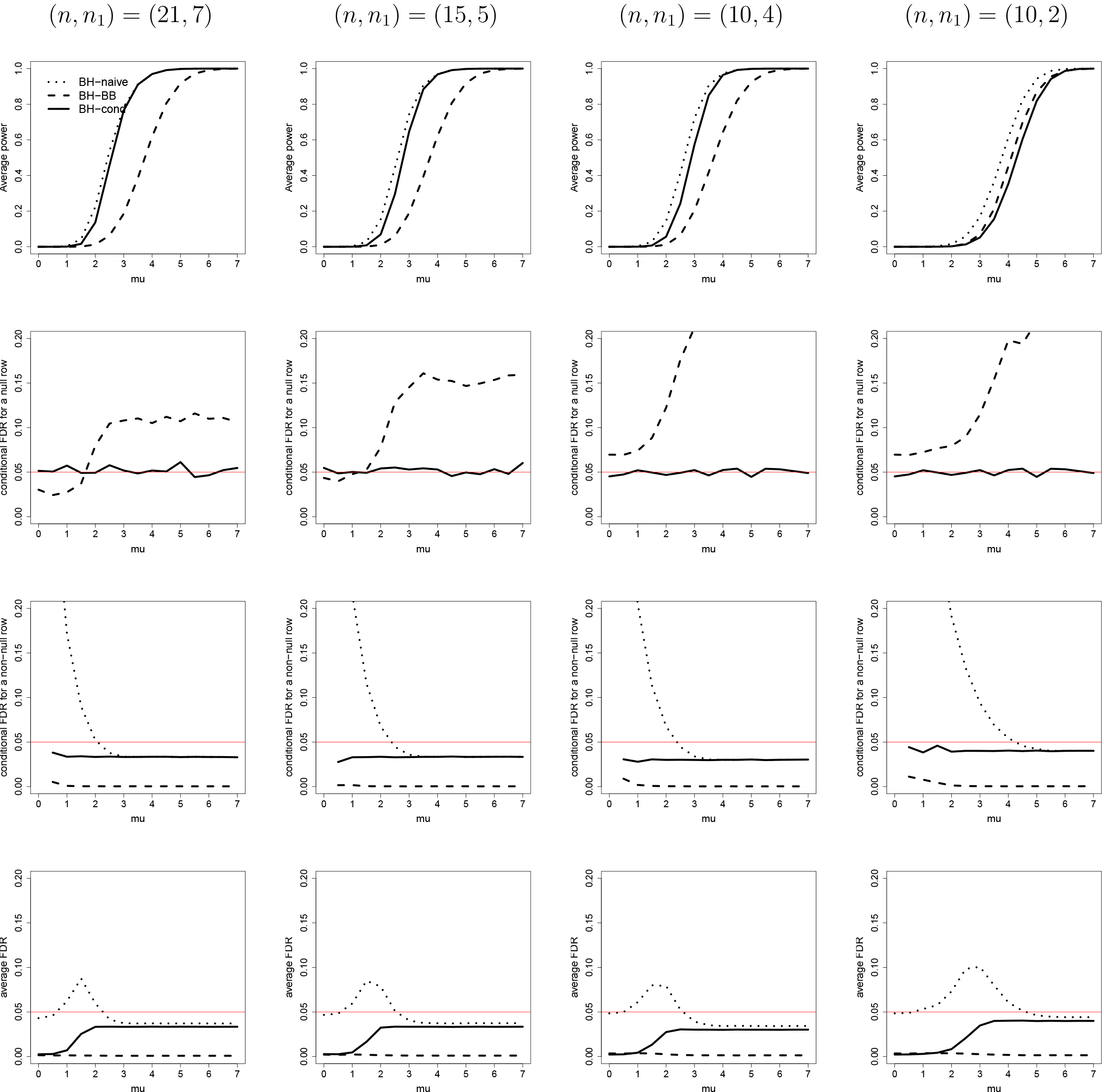
From left to right, variations in the total number of studies (*n*), and the number of studies with signal (ni) within the *m*_1_ = 10 rows that contained signal, out of a total of *m* = 1000 rows. From top to bottom: (1) average power, i.e., the expected fraction of discoveries among the true signals; (2) conditional FDR for a null row, i.e., a row that was selected despite the fact that all columns are null (the naive procedure does not appear because its value was above 0.80); (3) conditional FDR for a non-null row, i.e., a row that was correctly selected since it has signal in at least one column; and (4) average FDR. The power and error rates as a function of p (the signal strength in the non-null studies) of the three post selection analyses using the BH procedure: at level 0.05 on the original *p*-values (dotted line); at level |𝒮|0.05/*m* on the original *p*-values (dashed line); at level 0.05 on the conditional *p*-values in (3.4) (solid line). Estimated using 100000 datasets.

The results on the power show that the conditional approach is best when (1) many correctly selected rows are expected to contain signal in more than two columns; and (2) the number of rows selected out of the large number of rows examined is expected to be small (Figure 1). Note that in this simulation the average power is the same as the average power within a non-null row, and they are equal to the probability of discovering a true signal (i.e., rejecting a single non-null hypothesis), since we generated the signal (i.e., non-null test statistics) from the same distribution, and the same number of signals (i.e., non-nulls) in each of the *m*_1_ rows. Although the naive procedure should not be used due to its unacceptable inflation of false positives, we plot its power so it can serve as a benchmark for the power loss due to the necessary adjustment for selection. Examining the power of BH-BB and BH-cond, we see that the conditional approach is more powerful than the approach of Benjamini and Bogomolov (2014) when (*n*,*n*_1_) ∈ {(21, 7), (15, 5), (10, 4)}. The gain in power can be very large. For example, when μ = 3 the power difference between our approach and that of Benjamini and Bogomolov (2014) was greater than 40% for (*n*_1_,*n*) = (7, 21) and (*n*_1_,*n*) = (5,15) and about 30% for (n_1_,n) = (4,10). Our approach was slightly less powerful than the approach of Benjamini and Bogomolov (2014) for (*n*,*n*_1_) = (10, 2). Additional results are provided in Supplemental Table S1.

### 4.2 Results for the dependence setting

The results on the average and conditional FDR control show that the control of false positives was tightest using the conditional approach. The conditional approach controlled all the error measures (as expected) at the nominal 0.05 level when the row selection was by Bonferroni (i.e., the threshold for selection was independent of the global null *p*-values). When the row selection was by BH at level *q* = 0.2 on the global null *p*-values, the average FDR and the conditional FDR for a non-null row were below the nominal 0.05 level. However, the conditional FDR for a null row was slightly above the nominal 0.05. When the row selection was by BH at level *q* = 0.05 instead of q = 0.2, no inflation was observed (results not shown).

The approach of Benjamini and Bogomolov (2014) controlled at the nominal 0.05 level the average FDR/FWER (as expected), as well as the conditional FDR/FWER for a correctly selected row (i.e., a row that contains signal), but did not control the conditional FDR/FWER for an incorrectly selected row (i.e., a row that contains no signal). The naive approach exceeded the nominal 0.05 level in all error measures considered. Figure 2 shows the actual level of each error measures for the three post selection analyses procedures.

As in the independence setting, the results on the power show that the conditional approach is best when (*n*,*n*_1_) = (21, 7), but the approach of Benjamini and Bogomolov (2014) has slightly better power when (n, n_i_) = (10, 2) (Figure 2). Note that in this simulation the average power is different from the average power within a specific row, since the generated signal varied across rows.

**Figure 2:**
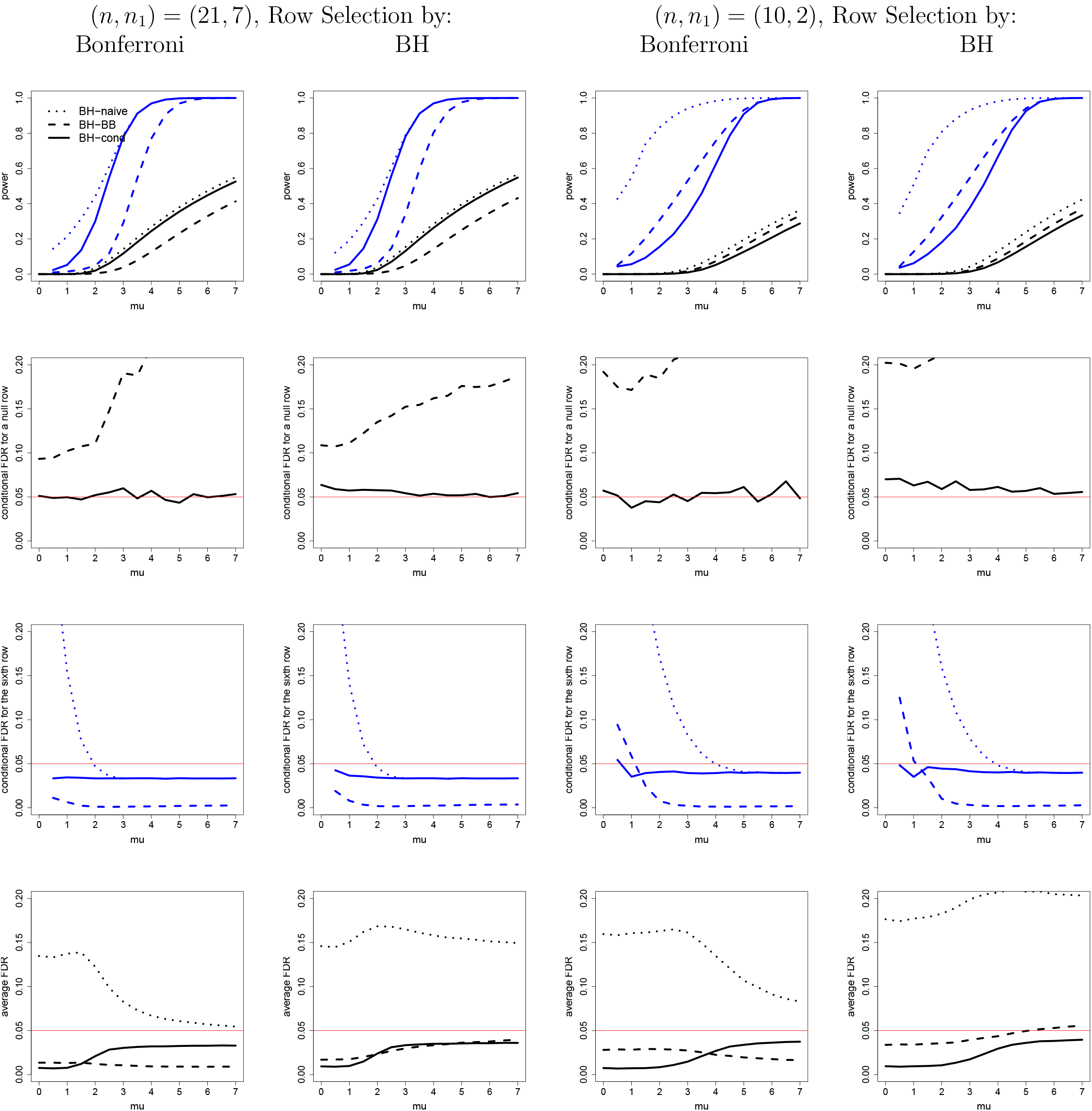
From left to right, *(n,n_i_) =* (21, 7) with row selection rule p_iG_ < 0.05/1100; *(n,n_i_) =* (21, 7) with row selection by BH at level q = 0.2 on {p_iG_, i = 1,…, 1100}; (n, *n*_i_) = (10,2) with row selection rule p_iG_ < 0.05/1100; and (n, ni) = (10, 2) with row selection by BH at level q = 0.2 on {p_iG_, i = 1,…, 1100}. Each study had 100 blocks of size 11 rows, and in n_i_ of the studies the first block contained signal. The dependence parameter was *p* = 0.7. From top to bottom: (1) average power (in black) and average power within the sixth row, which has the strongest signal in the block (in blue); (2) conditional FDR for a null row, i.e., a row that was selected despite the fact that all columns are null (the naive procedure does not appear because its value was above 0.80); (3) conditional FDR for the sixth row; and (4) average FDR. The power and error rates as a function of μ (the signal strength in the non-null studies) of the three post selection analyses using the BH procedure: at level 0.05 on the original *p*-values (dotted line); at level |S|0.05/m on the original *p*-values (dashed line); at level 0.05 on the conditional *p*-values in (3.4) (solid line). Estimated using 100000 datasets.

## 5 Cross-tissue eQTL analysis in The Cancer Genome Atlas (TCGA) Project

Expression quantitative trait loci (eQTLs) are genomic regions with genetic variants that influence the expression level of genes. Identifying eQTLs is important for understanding biological mechanisms that controls various normal physiological processes and their aberrations that can lead to complex diseases. Because gene regulation is tissue specific, eQTL analysis is most informative using relevant tissue samples, which suffers frequently from the lack of statistical power because of the small sample size for the tissue. It is, however, observed that some eQTL SNPs are predictive of gene expression levels across multiple tissues and identification of such eQTLs could be facilitated by aggregated analysis across tissue types. A number of studies (Rivas et al., (2015) and Li et al., (2016), among others) have reported results based on such cross-tissue eQTL analysis using the data from the Genotype-Tissue Expression (GTEx) project. In this section, we illustrate the post-selection procedure in an eQTL analysis using 17 tumor tissues in The Cancer Genome Atlas (TCGA) project (http://cancergenome.nih.gov/). We first performed an aggregated eQTL analysis across 17 tissue types to identify eQTL SNPs influencing the gene expression in at least one tissue type. For significant eQTLs, we performed post-selection inference to identify tissue types with the eQTL effect. We downloaded genotype and total gene expression data based on RNA sequencing from the TCGA website. The data quality control (QC) for genetic data and the normalization for the RNA-seq data are described in Supplementary Materials. After QC, 4,476 subjects with European ancestry and 19,285 genes were included for analysis. The sample size for each tissue type is reported in Supplementary Table S2. For the purpose of illustration, we performed cis-eQTL analysis (i.e., analyzing SNPs less than 1,000,000 base pairs from the target gene) using matrixeQTL (Shabalin, 2012), adjusting for sex, age and the top five principal component scores to eliminate the potential confounding due to population stratification. In total, we analyzed *m* = 7, 732, 750 SNP-gene pairs to identify cis-eQTLs. Because samples are independent across tissue types in TCGA, the *p*-values are independent across tissues.

For selection of eQTL SNPs (i.e., the rows), we combined *p*-values across the different tissue types (columns) by a Fisher style test suggested by Pearson, as described in Owen (2009): it runs the Fisher combining method for left-sided alternatives, and separately for right-sided alternatives, and takes the maximum of the two resulting statistics. The global null p-value is therefore

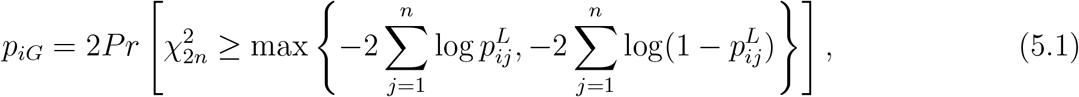

where 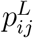 is the *p*-value when testing the left sided alternative for feature i in study j. This test has a strong preference for common directionality (i.e., it will have greater power than a test based on Fisher's combining method on two-sided *p*-values when the direction of the signal is consistent across tissues), while not requiring us to know the common direction. We identified 19,690 significant SNP-gene pairs using the global null p-value threshold 0.05/7, 732, 750 = 6.47 x 10^−9^ based on the Bonferroni correction for FWER control at the 0.05 level. For each of the 19,690 SNP-gene pair, we proceeded to post-selection inference to identify relevant tissue types using our conditional approach.

Table 1 demonstrates the post-selection analysis for three SNP-gene pairs, which differ in the number of identified tissue with signal in the post-selection inference. For the pair rs13066873- LARS2, the conditional *p*-values are identical to the original *p*-values, i.e., there is no cost for selection, since this pair would have been selected regardless of the realized p-value in a single tissue, when conditioning on all the other tissue *p*-values for this pair. From the BH adjusted *p*-values provided in column 10, we see that with the conditional approach 15 tissues are discovered which have adjusted *p*-values at most 0.05, but with the approach of Benjamini and Bogomolov (2014) only the tissue LUAD is discovered, i.e., the single tissue with BH-adjusted p-value at most 0.05 x 19, 690/7, 732, 750 = 0.00013. For the pair rs1437891-ASNSD1 in columns 5-7, with the conditional approach six tissues are discovered, and these tissues have conditional *p*-values larger than the original *p*-values. With the approach of Benjamini and Bogomolov (2014) no discoveries are made. For the pair rs7977641-GALNT9 in columns 2-4, no discoveries are made in either method.

**Table 1:** The original two-sided *p*-values, conditional two-sided *p*-values, and BH-adjusted conditional two-sided *p*-values for each tissue, for three eQTL SNPs that differ in the number of post selection discoveries: rs10896016-CTSW identified no tissues (columns 2-4); rs1437891-ASNSD1 identified 6 tissues (columns 5-7); rs13066873-LARS2 identified 15 tissues (columns 8-10). Significant discoveries at the 0.05 FDR nominal level are in bold for the BH-adjusted conditional *p*-values. The global null *p*-value for each SNP-gene pair are provided in the last row.

| | rs10896016-CTSW $p$-values | | | rs1437891-ASNSD1 $p$-values | | | rs13066873-LARS2 $p$-values | | |
| --- | --- | --- | --- | --- | --- | --- | --- | --- | --- |
|  | original | conditional | BH-adjusted | original | conditional | BH-adjusted | original | conditional | BH-adjusted |
| BLCA | 1.26e-02 | 2.95e-01 | 3.86e-01 | 4.55e-01 | 4.55e-01 | 6.45e-01 | 1.99e-03 | 1.99e-03 | **4.84e-03** |
| BRCA | 7.33e-01 | 7.33e-01 | 8.30e-01 | 2.98e-04 | 8.04e-03 | **2.28e-02** | 2.60e-04 | 2.60e-04 | **1.47e-03** |
| COAD | 2.66e-01 | 2.95e-01 | 3.86e-01 | 2.31e-03 | 2.31e-03 | **2.28e-02** | 9.93e-04 | 9.93e-04 | **3.62e-03** |
| GBM | 3.61e-01 | 2.95e-01 | 3.86e-01 | 9.02e-01 | 9.02e-01 | 9.02e-01 | 7.16e-03 | 7.16e-03 | **1.35e-02** |
| HNSC | 9.22e-01 | 9.22e-01 | 9.80e-01 | 5.47e-01 | 5.47e-01 | 6.64e-01 | 5.44e-01 | 5.44e-01 | 5.44e-01 |
| KIRC | 7.43e-03 | 2.95e-01 | 3.86e-01 | 2.56e-07 | 8.04e-03 | **2.28e-02** | 1.36e-02 | 1.36e-02 | **1.78e-02** |
| KIRP | 9.96e-01 | 9.96e-01 | 9.96e-01 | 5.20e-01 | 5.20e-01 | 6.64e-01 | 8.34e-03 | 8.34e-03 | **1.42e-02** |
| LAML | 2.35e-02 | 2.95e-01 | 3.86e-01 | 7.78e-01 | 7.78e-01 | 8.27e-01 | 3.45e-03 | 3.45e-03 | **7.33e-03** |
| LGG | 1.40e-01 | 2.95e-01 | 3.86e-01 | 4.54e-05 | 8.04e-03 | **2.28e-02** | 1.07e-03 | 1.07e-03 | **3.62e-03** |
| LIHC | 1.57e-02 | 2.95e-01 | 3.86e-01 | 3.44e-01 | 3.44e-01 | 6.45e-01 | 1.01e-02 | 1.01e-02 | **1.43e-02** |
| LUAD | 3.58e-05 | 2.95e-01 | 3.86e-01 | 7.79e-04 | 8.04e-03 | **2.28e-02** | 1.24e-07 | 1.24e-07 | **2.11e-06** |
| LUSC | 1.29e-01 | 2.95e-01 | 3.86e-01 | 3.03e-01 | 3.03e-01 | 6.45e-01 | 4.07e-02 | 4.07e-02 | **4.83e-02** |
| OV | 6.66e-02 | 2.95e-01 | 3.86e-01 | 1.63e-01 | 1.63e-01 | 3.95e-01 | 9.61e-03 | 9.61e-03 | **1.43e-02** |
| PAAD | 2.57e-01 | 2.57e-01 | 3.86e-01 | 6.42e-01 | 6.42e-01 | 7.27e-01 | 4.26e-02 | 4.26e-02 | **4.83e-02** |
| PRAD | 1.41e-01 | 2.95e-01 | 3.86e-01 | 4.95e-03 | 8.04e-03 | **2.28e-02** | 6.41e-02 | 6.41e-02 | 6.81e-02 |
| SKCM | 1.58e-02 | 2.95e-01 | 3.86e-01 | 4.15e-01 | 4.15e-01 | 6.45e-01 | 1.83e-04 | 1.83e-04 | **1.47e-03** |
| UCEC | 5.92e-01 | 5.92e-01 | 7.19e-01 | 4.29e-01 | 4.29e-01 | 6.45e-01 | 1.67e-03 | 1.67e-03 | **4.73e-03** |
| Global null $p$-value | $3 \times 10^{-9}$ | | | $2 \times 10^{-10}$ | | | $< 10^{-20}$ | | |

Using the BH procedure on selected rows, the median number of tissue discoveries was 6, with an inter-quartile range (IQR) of [4,8]. For comparison, we also applied the approach of Benjamini and Bogomolov (2014), which made far fewer discoveries: the median number of discoveries was 1, with IQR [0,2]. The conditional approach results in many more discoveries than the approach of Benjamini and Bogomolov (2014) for two reasons. First, because the number of selected SNP-expression pairs is far smaller than the number originally examined, 19,690/7,732,750=0.0025, and the approach of Benjamini and Bogomolov (2014) is more conservative the smaller this ratio is. Second, because for many SNP-expression pairs there are at least two tissues which are highly significant, thus the conditional *p*-values coincide with the original *p*-values, and there was essentially no cost for selection within these pairs.

An *R* implementation of the conditional *p*-value computation after selection by thresholding the global null *p*-values computed using (5.1) or (3.1) is available upon request from the first author (and will soon become available as a Bioconductor package).

### 5.1 Valid inference within the selected most significant SNP-expression pair in a gene

For a target gene, there might be multiple SNPs in the cis region that achieve the genome-wide significance. Most likely, these SNPs are in strong linkage disequilibrium (LD) and represent one eQTL. To avoid reporting redundant eQTLs, one natural step is to choose the SNP with the smallest global null *p*-value and perform post-selection inference to identify relevant tissue types. However, the post-selection inference may suffer from high false positive rate if this second selection is not appropriately accounted for. Related simulation results are provided in Supplemental Figure S2. Thus, we propose a simple modification to our post-selection inference method to account for the second selection. We denote the recommended procedure for FDR control by BH-cond-MT, where MT stands for the additional Multiple Tests (of the SNPs that passed the first selection threshold at the aggregate level in the gene) that we need to correct for, after computing the conditional *p*-values.

#### Procedure 5.1.

*BH-cond-MT post-selection procedure for conditional FDR control at level α*:

1. *Select all SNP-expression pairs that reach the genome-wide significance threshold h(q, m) (e.g., h(q, m) = q/m if the Bonferroni correction is used for FWER control at level q on the global null hypotheses), 𝒮 = {i: p_iG_ < h(q, m)}*.
2. *For each gene k with at least one selected SNP, select the most significant SNP in gene k, i_k_ =* arg min_{*i:i*∈𝒮*i in gene k*_}*p_iG_*. *Let R_k_* = |{*i*: *i* ∈ 𝒮,*i in gene k*}| *be the number of SNPs selected in gene k*.
3. *Compute the conditional p-values as in (3.3) for row i_k_*.
4. *At level α/R_k_ on the conditional p-values in row i_k_, apply the BH procedure.*

For conditional FWER control, apply Bonferroni-Holm instead of BH in step 4 of Procedure 5.1. In Supplemental Figure S3 we show that the procedure controls the FDR for dependencies across SNPs similar to those that arise in GWAS datasets within each study due to LD, and in Appendix D we formally prove that the conditional FDR is controlled when the rows are independent.

The impact of the second selection is illustrated in Table 2 for gene KIAA0141. Four SNPs (rs351260, rs164515, rs164084, rs164075) were identified for KIAA0141. Out of the four SNPs, rs351260 had the strongest overall association and was selected for post-selection inference to identify relevant tissues. Without accounting for the second selection, 14 tissue types were determined as significant based on the BH-adjusted conditional *p*-value < 0.05. After accounting for the second selection, 9 tissue types were counted as significant based on the new threshold BH-adjusted conditional *p*-value*<* 0.05/4 = 0.0125.

**Table 2:** The original *p*-values for the four SNPs for gene KIAA0141 that passed the genome-wide selection threshold (columns 3 to 6), as well as the BH adjusted conditional *p*-values for the most significant SNP (column 6). In bold the adjusted *p*-values <, i.e., the significant discoveres using Procedure 5.1 with *α* = 0.05. Underlined are the adjusted *p*-values in (0.05/4,0.05], i.e., discoveries using BH-cond that are not discoveries using Procedure 5.1 with *α* = 0.05. The conditional *p*-values were identical to the unconditional *p*-values in all four SNPs (since for each tissue *j*, the SNP would have been selected regardless of the value of the *j*th *p*-value). The global null test-statistic for each SNP is provided in the last row (the corresponding *p*-value was effectively zero).

| tissue type | $p$-value<br>rs164515 | $p$-value<br>rs164084 | $p$-value<br>rs164075 | $p$-value<br>rs351260 | BH adjusted $p$-value<br>rs351260 |
| --- | --- | --- | --- | --- | --- |
| BLCA | 1.94e-02 | 1.32e-01 | 1.51e-01 | 2.79e-03 | **5.93e-03** |
| BRCA | 1.10e-02 | 2.11e-02 | 1.85e-02 | 2.79e-04 | **7.92e-04** |
| COAD | 3.48e-01 | 3.01e-01 | 3.01e-01 | 3.67e-02 | <u>4.45e-02</u> |
| GBM | 8.69e-02 | 5.61e-01 | 5.91e-01 | 7.92e-02 | 8.97e-02 |
| HNSC | 3.93e-01 | 2.16e-01 | 2.03e-01 | 5.69e-01 | 5.69e-01 |
| KIRC | 5.13e-07 | 1.87e-05 | 7.75e-06 | 1.56e-07 | **1.33e-06** |
| KIRP | 1.25e-02 | 3.37e-02 | 3.05e-02 | 3.00e-02 | <u>4.03e-02</u> |
| LAML | 1.33e-01 | 6.54e-02 | 9.01e-02 | 5.66e-03 | **1.07e-02** |
| LGG | 1.14e-04 | 7.68e-02 | 8.90e-02 | 6.39e-07 | **3.62e-06** |
| LIHC | 4.01e-02 | 1.21e-01 | 7.87e-02 | 3.08e-02 | <u>4.03e-02</u> |
| LUAD | 3.31e-03 | 1.42e-02 | 1.02e-02 | 3.72e-04 | **9.04e-04** |
| LUSC | 1.71e-07 | 1.49e-05 | 3.32e-05 | 1.18e-07 | **1.33e-06** |
| OV | 5.45e-04 | 1.11e-01 | 7.88e-02 | 2.68e-04 | **7.92e-04** |
| PAAD | 1.79e-01 | 9.42e-02 | 1.40e-01 | 4.32e-01 | 4.59e-01 |
| PRAD | 2.58e-03 | 2.86e-03 | 2.11e-03 | 7.32e-05 | **3.11e-04** |
| SKCM | 5.95e-02 | 8.55e-01 | 6.76e-01 | 8.33e-03 | <u>1.42e-02</u> |
| UCEC | 3.14e-02 | 3.58e-02 | 2.25e-02 | 2.16e-02 | <u>3.34e-02</u> |
| global null test-statistic | 343.7 | 361.6 | 346.0 | 396.8 |  |

Finally, we compared the performance of four methods based on the whole eQTL analysis: BH-naive, BH-cond and BH-cond-MT and BH-BB (Table 3). Note that BH-naive and BH-cond do not account for second selection and thus their error rates are likely to be inflated. Consistent with simulation studies, BH-BB detected the smallest number of signals because of lower power.

**Table 3:** By each post-selection method, the number of genes with at least *x* discoveries of tissues, for *x* = 0,1,…, 14. The first row represents the genes with at least 0 discoveries, i.e., the number of selected genes. The theoretically valid methods, which control their respective error rates, are BH-BB (column 5), as well as the methods that adjust for the selection of a single row per gene but use the conditional *p*-values, BH-cond-MT (column 4), described in Procedure 5.1.

| minimum #<br>of discoveries | BH-naive | BH-cond | BH-cond-MT | BH-BB |
| --- | --- | --- | --- | --- |
| 0 | 2235 | 2235 | 2235 | 2235 |
| 1 | 2235 | 2003 | 1936 | 1309 |
| 2 | 2200 | 1980 | 1857 | 678 |
| 3 | 2105 | 1938 | 1653 | 352 |
| 4 | 1891 | 1808 | 1352 | 160 |
| 5 | 1642 | 1615 | 983 | 59 |
| 6 | 1348 | 1333 | 677 | 7 |
| 7 | 1081 | 1081 | 447 | 2 |
| 8 | 832 | 832 | 266 | 0 |
| 9 | 604 | 604 | 132 | 0 |
| 10 | 375 | 375 | 50 | 0 |
| 11 | 195 | 195 | 15 | 0 |
| 12 | 84 | 84 | 8 | 0 |
| 13 | 21 | 21 | 0 | 0 |
| 14 | 1 | 1 | 0 | 0 |

For example, among the 2235 selected SNPs, at least two tissue discoveries were made in 1857 of the genes by BH-cond-MT and only in 678 of the genes by BH-BB. For the BH procedure based on conditional *p*-values, accounting for second selection noticeably reduced the significant findings. This was expected, since the number of SNPs discovered per gene is typically greater than one: out of the 19690 pairs that reached genome-wide significance, there were 2235 unique genes, and the number of SNPs per gene varied between 1 and 76, with a median number of 5.

## 6 Discussion

Results from both simulation studies and data analysis highlight the potential of the proposed method for valid and powerful hypothesis testing for detection of signals at the level of the finer units following selection of broader units using aggregate level test-statistics. Although the method is not as general as its existing competitor (Benjamini and Bogomolov, 2014), it can handle an important class of scenarios that involve independence of the primary test-statistics across columns, a practical context of which is demonstrated through the application involving cross-tissue eQTL analysis in the rich TCGA dataset. The superior power of the proposed procedure over that of Benjamini and Bogomolov (2014) in this particular setting implies that a general error-controlling method may not be very powerful for specific applications. Thus, substantial scope for future research exist for development of other powerful procedures tailored towards specific important application settings following the general principles we lay out.

If the columns are dependent, it is an open question how to compute valid conditional *p*-values. In this work we relied on the fact that the null distribution for a unit-level test statistic is known when we condition on the selection event and on all the other *p*-values in its row. When the columns are dependent, the null distribution after conditioning on the selection event and on all other *p*-values in the row may still depend on unknown parameters. The approach of Benjamini and Bogomolov (2014) remains valid in this case, since it is not sensitive to dependence across columns, as long as the within row multiple testing procedure controls the desired error rate for the dependence. In applications where the dependency across columns is approximately known, it may be possible to compute the conditional *p*-values and carry on the post-selection inference as we suggest in this paper. We plan to investigate the usefulness of this approach for specific applications in future work.

Other post-selection analyses may be of interest. For example, estimation of the fraction of columns containing signal within each selected row. Such estimates can be useful in separating the selected rows where there is signal in most columns, from the rows driven by very few (one or two) columns only that contain signal. Another example is the estimation of a linear combination of the effect sizes. A conditional approach can be useful for these post-selection estimation problems.

## A Proof of Corollary 3.1

*Proof*. We let the first hypothesis be a true null, and reorder and relabel the columns so *p_2_* ≤ … ≤ *p_n_*. Let *J* = argmax_2_≤_k_≤_n_{*p*_k_ ≤ *αk/n}*. If no such *k* exists then *J* =1. Note that *R* ≤ *J*, since the number of *p_2_,…, p_n_* that does not exceed *αk/n* is at most k − 2 for k > J. Since *p_1_^′^* ≥ *p_j_* there are at most k − 2 of the *p*_j_^′^s that do not exceed *αk/n*. Even if *p*_1_^′^ ≤ *αk/n*, the kth largest from among *p*_1_^′^,…,*p*_n_^′^ exceeds *αk/n*. It follows that if *p_1_^′^ > αJ/n* then *I* = 0.

If *p_1_^′^ ≤ αJ/n* and *b_1_* = 1, for *r ≤ J*: *p_r_^′^* is either equal to *p_r_^′^* and therefore *p_r_^′^ ≤ αJ/n*, or 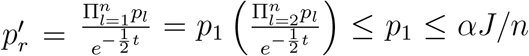.If *p_1_^′^ ≤ αJ/n* and *b_1_ <* 1, for *r ≤ J*: since 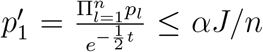,then *p_r_^′^ ≤ αJ/n*. Therefore, if *p*_r_^′^ ≤ *αJ/n* then *p*_r_^′^ ≤ αJ/n for *r ≤ J*, i.e., *R* = *J*.

Therefore,

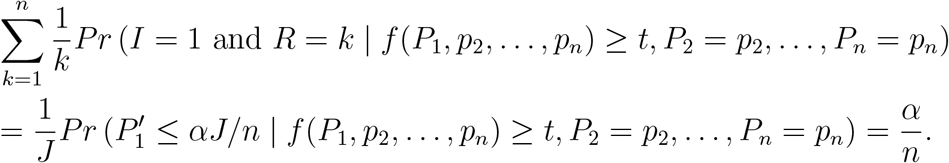

## B Proof of Theorem 3.2

Since the selection rule is simple and we are conditioning on *P^(i)^*, there will be a unique number of rows that are rejected, *k* along with *i*, which depends on *P^(i)^*. Let *C*_+_ be the possible values of *C_i_* ∈{*I*[*V_i_*> 0],*V_i_*/*max*{*R_i_*, 1}}.

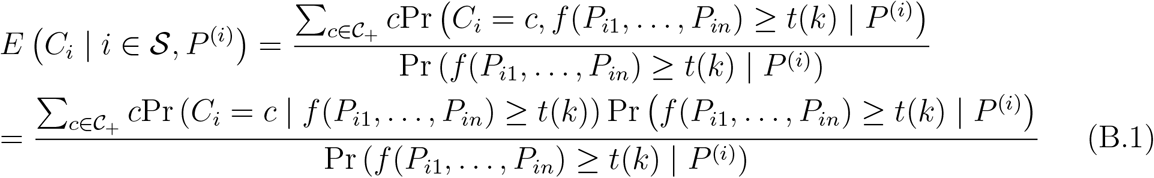

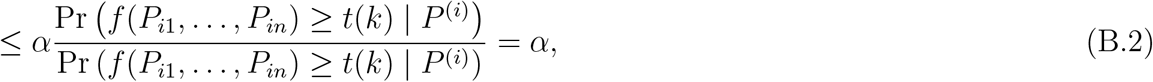
where equality (B.1) follows since the rows are independent. Inequality (B.2) follows since level-α conditional inference controls level-α conditional error for selection thresholds that do not depend on **p*_i1_, …, *p*_in_*, as proved in Section 3.1.2.

## C Proof of Theorem 3.3

*Proof*. Since *E*(*C_i_* | | *i* ∈ *𝒮, P*^(*i*)^) ≤ *α*, it follows that

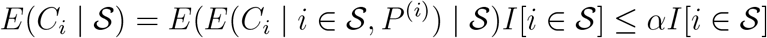

The result is immediate:

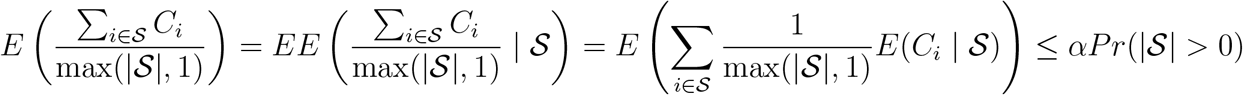

## D Proof of conditional FDR control for Procedure 5.1

Procedure 5.1 controls the conditional FDR when the rows are independent, as formally stated in the following theorem.

### Theorem D.1.

*Assume the global null p-values in gene k are independent. If R_k_ of the SNPs in gene k have global null p-values at most h*(*q*,*m*), *then the Bonferroni-Holm/BH procedure at level α/R_k_ on* 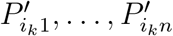, *where i_k_* = arg min_{*i*:*i*∈*𝒮 i in gene k*}_ *p_iG_, controls the conditional FWER/FDR at level* ≤*α*.

*Proof*. Relabel the rows so that the first *m_k_* rows are the SNPs for gene k.Let 𝒮_k_ be the selection status for the SNPs in gene k, i.e., 𝒮_k_ = {I*[*P*_IG_ ≤ h(q,m)],i* in gene *k*}.

Let 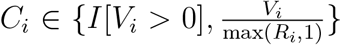. The procedure guarantees conditional error control for each SNP i in gene *k* at level *α*/*R_k_* if the rows are independent:

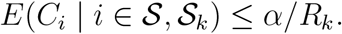

It therefore follows that the conditional error is controlled also for the row with the smallest global null p-value:

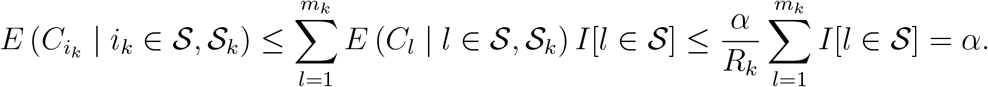

## Notes

2 Study supported by the National Cancer Institute Intramural Research Program.

